# Archaeal cell envelopes selectively activate innate immunity through TLR2 in contrasts with multi-receptor bacterial sensing

**DOI:** 10.64898/2026.10.03.756221

**Authors:** Camille Martin-Gallausiaux, Charles Grosjean, Brittany Baker, Catherine Werts, Guillaume Borrel, Simonetta Gribaldo

## Abstract

The cell envelope of bacteria and fungi is a major source of microbial-associated molecular patterns (MAMPs), which are recognised by multiple host pattern recognition receptors (PRRs) such as TLRs and NODs. The key role of this recognition in protecting against infection and regulating diverse physiological processes has been studied for decades. In contrast, although the archaea are growingly recognised as an important component of the animal and human microbiome, their interaction with the innate immune system has been largely understudied. Here, we asked whether the archaeal cell envelope acts as MAMP. We selected 15 archaeal species from different environments, including the human microbiome, and harbouring diverse envelope characteristics, and we tested their ability to activate innate immune responses in human and murine immune cells. We first show that archaeal cells activate the innate immune transcription factor NF-kB and induce IL-6 secretion. Using whole archaeal cells, supernatants, and purified envelope components, we then find that none of the main PRR receptors which recognise the bacterial envelope (TLR4, TLR5, NOD1 and NOD2) are involved in this activation. We only detected activation of TLR2, regardless of the environmental origins of the archaea tested, although with overall lower activity compared to bacteria. We further demonstrate that different TLR2 heterodimers are involved, depending on the species, although the archaeal agonist remains unknown. Altogether, our results highlight a surprising paucity of interaction of archaeal cell envelope components with host PPRs, in contrast to bacteria, and open new avenues of research toward further investigation of archaea/host interplay.

## Introduction

In 1989, Charles Janeway proposed the concept of “pattern recognition model” to integrate unexplained observations of both innate and adaptive immune responses in mammals. He hypothesised that eukaryotes should detect all members of a given group of microorganisms due to conserved microbial products recognised by a limited repertoire of non-variable receptors^1^. A decade later, the membrane Toll-like receptors (TLRs) were discovered, as well as other pattern recognition receptors (PRRs), such as the cytosolic NODs, and some of their ligands, generally referred to as microbial-associated molecular patterns (MAMPs)^2^.

MAMPs share several commons features: they are widely distributed, abundant, essential for the viability or fitness of microorganisms in their ecological niches, and absent from the host. MAMPs can be divided in two groups: nucleic acids-derived and cell envelope-derived^3^. The most studied cell envelope-derived MAMPs include bacterial lipopolysaccharide (LPS) recognised by TLR4, lipoproteins recognised by TLR2, flagellins recognised by TLR5, and fragments of peptidoglycan (PG) recognised by NOD1 and NOD2^3–5^. The fungal cell envelope also provides various MAMPs such as chitin, β-glucans, zymosan and mannans, although their direct interactions with TLRs and NODs are much less characterised^6^.

In contrast, the MAMPs-PRRs landscape of the cell envelope of Archaea, one of the three domains of life, is largely unknown. Archaea are present in all environments, including oceans, soil, and extreme environments^7,8^. Some species are associated with a large variety of hosts including insects, birds, fishes, and humans^9,10^. Archaea have co-evolved for millions of years with humans as commensals and are part of the oral, gastrointestinal, and skin microbiota^11,12^. In the human gut, the four main species of archaea are all strict anaerobic methanogens: *Methanobrevibacter smithii, Methanobrevibacter intestini, Methanosphaera stadtmanae* all three belonging to the Methanobacteriales*, and Methanomethylophilus alvi* (Methanomassiliicoccales), with a predominance of the first two species^11^. The dominant species in the oral cavity is *Methanobrevibacter oralis*^11^. Archaea belonging to several lineages of methanogens, halophiles, and ammonia oxidizers, have also been detected on other mucosa, as well as on the skin^11,13^. Very few data exist on the role of intestinal archaea in human health and disease, but they have been shown to be decreased in Crohn’s disease and obesity and increased in irritable bowel syndrome with constipation^14,15^. Moreover, although no archaeal pathogen has been reported in humans to date, archaea have been linked with infections, notably by a correlation of the relative abundance of *M. oralis* and periodontitis severity^12,16,17^.

A distinctive feature of archaea with respect to bacteria lies in their cell envelopes^18,19^. These are mostly constituted of a single lipid membrane, organised as a bilayer or a membrane-spanning monolayer. Archaeal membrane lipids are composed of isoprenoids chains bound to a glycerol backbone via ether linkage as opposed to bacterial and eukaryotic membrane lipids which are composed of fatty acid chains bound by ester-linkages to the glycerol moiety ^20^. In the absence of an outer membrane as in diderm bacteria, archaea do not have LPS. The outermost surface of archaeal cells is often protected by an S-layer, a proteinaceous sheath, and/or a polysaccharide such as methanochondroitin^18^. Moreover, archaea generally lack peptidoglycan, except for Methanopyrales and Methanobacteriales which possess cell walls containing pseudomurein (archaeal PG). Archaeal PG is chemically distinct from that of bacteria and notably does not contain MurNAc^21,22^. Archaea also express various surface appendages including pili, fimbriae, cannulae, or hami^18^. Finally, archaeal motility is based on a flagellum that is not homologous to its bacterial counterpart, but rather resembles to the bacterial type IV pilus, and is therefore called archaellum^23^.

Given that archaea, similarly to bacteria, have co-evolved with mammals for millions of years^10^, it could be expected that their cell envelopes also produce MAMPs recognised by PRRs. Previous work has investigated the effect of *M. smithii* and *M. stadtmanae* whole cells in monocytes and epithelial cells and found that only *M. stadtmanae* induces cytokine secretion but no activity on TLRs and NODs^24^. A second study found that *Methanomassiliicoccus luminyensis* whole cells also induces cytokine secretion^25^. However, these studies did not specifically test different components of the archaeal cell envelope, nor a wide diversity of representatives of this domain.

Here, we carried out a thorough analysis of whether archaeal cell envelope-derived molecules function as MAMPs for TLR2-4-5 and NOD1-2. Our results demonstrate that, although these PRRs have evolved to detect multiple MAMPs derived from the bacterial envelope, they are surprisingly barely involved in recognition of the archaeal cell envelope, irrespective of their niches. We discuss the implications of our findings and the perspectives ahead.

## Results

### 1. Archaea from diverse environmental niches induce NF-kB

We selected 15 cultivable species of archaea covering a wide taxonomic diversity of this domain (Figure 1A). We included representatives of the major host-associated archaea, under the hypothesis that they are the most likely to be detected by PRRs. Archaea living in various environments were also selected, with the assumption that they may show different patterns in the absence of fitness pressure by the host. Moreover, these species cover a wide diversity of cell envelope structures: archaeal PG, archaeal flagella, and S-layer, all potential candidates for being MAMPs (Figure 1A).

**Figure 1.**
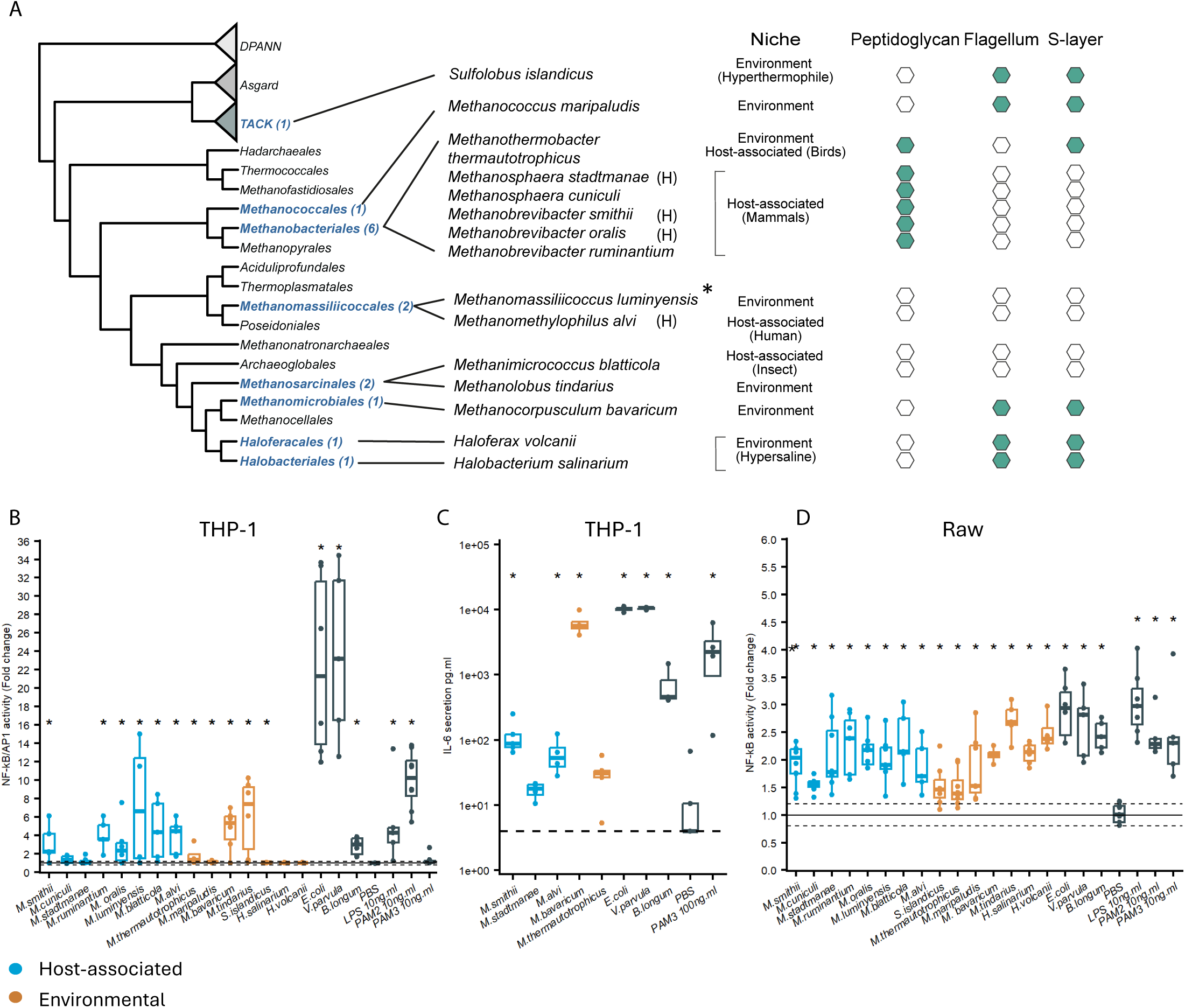
Diverse archaea activate human and murine PRRs. **A**) Schematic phylogeny of Archaea showing the distribution of the selected species within their corresponding clades (in blue), along with their habitat and cell envelope characteristics. Human-associated species are indicated by an H symbol. *M. luminyensis was isolated from human faeces but is mainly recovered from environmental samples. **B**) NF-kB/AP1 activation level after 24h exposure to archaeal cells on THP-1. **C**) secretion of IL-6 after 24h exposure of archaeal cells on THP-1 cells. **D**) NF-kB/AP1 activation level after 24h exposure to archaeal cells on Raw cells. NF-KB activation results are expressed as fold change compared to PBS treated cells, represented as the median ± quartiles (N ≥ 3). *p value < 0.05 by Wilcoxon rank test. Black: gram-negative controls Escherichia coli and Veillonella parvula; gram-positive Bifidobacterium longum. Additional controls are LPS (TLR4 ligand), and PAM2-3 (TLR2 ligands).

We first tested if the 15 species trigger the activation of innate immunity in human and mouse immune cells. We used the human monocyte reporter cell line (THP-1 Xblue NF-kB/AP1), as it expresses most PRRs and is highly sensitive to microbial MAMPs. Our results showed that most archaea activated these innate immune transcription factors, although the response was lower compared to the gram-negative bacterial controls (Figure 1B). The exposure of THP-1 to archaeal cells induced not only NF-kB activation but also the release of IL-6 cytokines, although the IL6 response for *M. bavaricum* exceeded the NF-kB response (Figure 1C). We also showed that NF-kB is quite similarly activated by archaea in human monocytes and in the NF-kB RAW murine macrophages cell line (Figure 1D). The only exceptions are halophilic archaea (*H. salinarum* and *H. volcanii*) which induce NF-kB activation only in murine cells (Figure 1B and C). However, this result must be taken with caution as these archaea were fixed prior to the assays to prevent cell lysis due to osmotic shock, which could induce a technical bias.

These results indicate that archaea induce innate immunity. To identify the PPRs involved in this recognition we used HEK reporter cells. We first confirmed that there was no background activity of archaea on the control reporter line (HEK Null), which express low levels of TLR3, TLR5 and NOD1^26^ (Supplementary Figure 1A), and similarly no response was observed on TLR4 overexpressing HEK cells (Supplementary Figure 1B).

We therefore proceeded to test the ability of the selected archaea to activate TLR2-4-5 and NOD1-2, by including supernatants and specific envelope elements based on knowledge of bacterial MAMPs.

### 2. TLR5 does not recognise archaea

TLR5 is a receptor with low flexibility in terms of ligands, and it specifically recognises the D0 and D1 domains of the bacterial flagellin^27^. Using HEK reporter cells overexpressing TLR5, we tested if this receptor could also detect archaeal cells, both flagellated and non-flagellated, as well as culture supernatants.

While both the flagellated *E. coli* control, and the purified bacterial flagellin activate human TLR5, no activation was detected upon stimulation with archaeal cells, whether they are flagellated or not (Figure 2A). Regarding archaeal supernatants, the only relevant activation compared to the control media concerns *M. bavaricum*, which showed a consistent albeit very low response (Figure 2B). As human and murine TLR5 receptors are known to differ in terms of ligand affinity^28^, we tested if archaeal molecules in the supernatants are recognised by the murine TLR5, however no activity was observed (Figure 2C).

**Figure 2.**
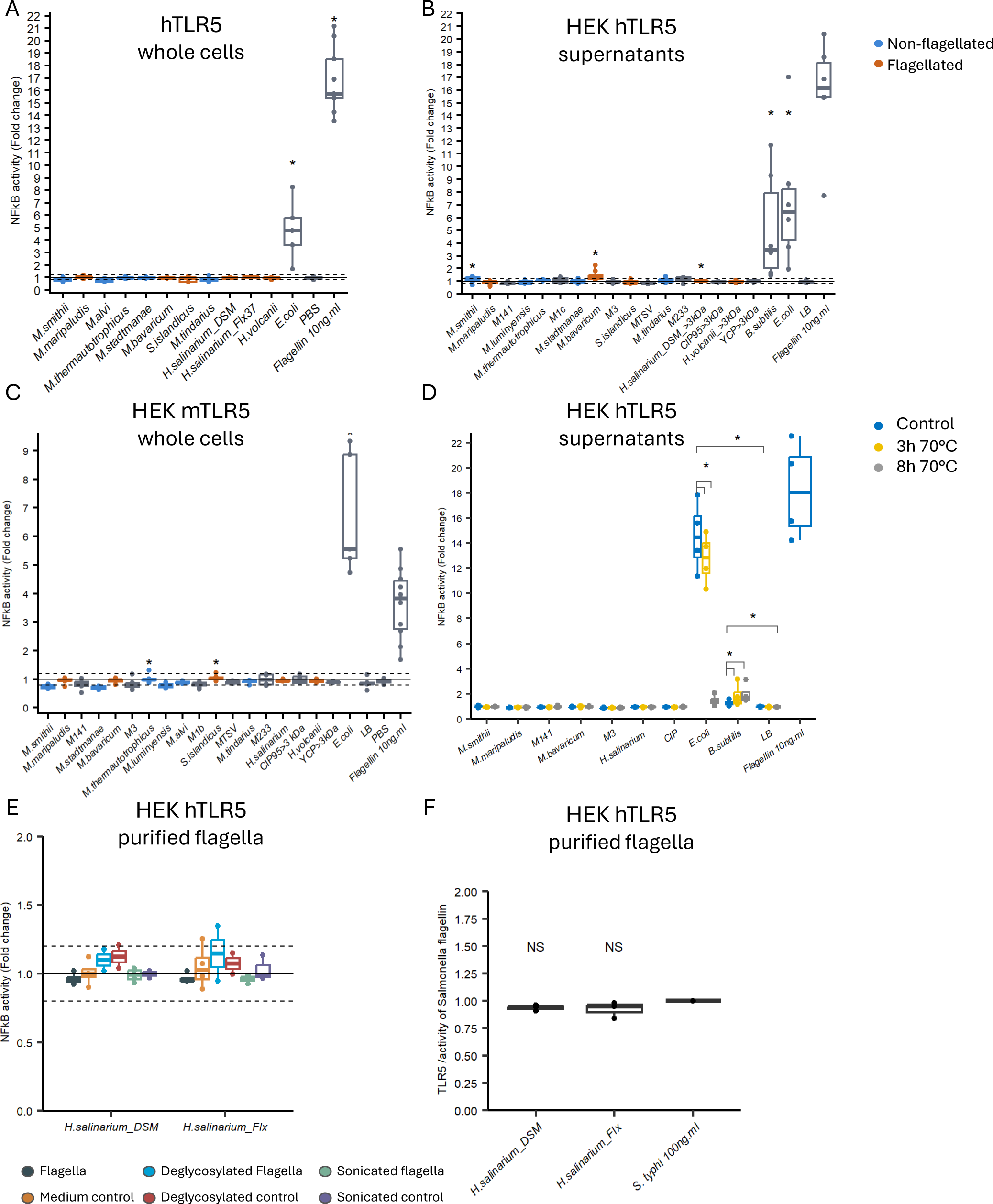
Archaeal flagella do not act as MAMPs for TLR5. NF-kB activation of hTLR5 after exposure to **A**) archaeal cells and **B**) supernatants and control bacteria. **C**) NF-kB activation of murine TLR5 after exposure to archaeal supernatants. **D**) NF-kB activation of hTLR5 after exposure to heat-treated supernatants. **E**) NF-kB activation of hTLR5 cells after exposure to extracted H. salinarium flagella, with or without deglycosylation and sonication treatments. Boxplots 2A-E show the median ± 1^st^ and 4^th^ quartiles. **F**) NF-kB/AP1 hTLR5 activation after exposure to purified archaeal flagella for 2h before treatment with S. Typhi flagella. NF-kB/AP1 is expressed as ratio of S. Typhi activity. NF-KB activation results are expressed fold change compared to the corresponding control media or PBS.

We investigated if the absence of recognition by TLR5 could be due to archaeal flagella polymerisation that might masks the TLR5 binding sites as described for bacterial flagellin^29^. However, depolymerisation of archaeal supernatant by prolonged heating did not result in higher TLR5 activation (Figure 2D). Glycosylation of the bacterial flagellum has been shown to be able to interfere with TLR5 recognition^30–32^. Archaeal flagella can also be glycosylated, especially in halophiles^33,34^. We therefore purified the flagella of *H. salinarium* (Supplementary Figure 2A and B), removed the N– and O-type glycosylation following published protocols^35,36^, and separated flagellar bundles by sonication. None of the purified archaeal flagella, either with or without deglycosylation and sonication, triggered TLR5 response (Figure 2E). We also asked if archaeal flagella could compete with or inhibit the activation of bacterial flagellin on TLR5. Antagonistic effects have in fact been described with the evasive flagella of α and ε Proteobacteria^37^ or the silent flagella of *Roseburia*^38^. However, pre-incubation with purified archaeal flagella prior to treatment with purified flagellin of *S. abortus* induced no reduction of the TLR5 response (Figure 2E).

We could not identify any D0 and D1 protein motifs in the proteomes of the 15 archaeal species, indicating that the lack of activation is not due to masking or competition for TLR5 binding sites.

### 3. NOD receptors do not sense archaeal PG

NOD1 and NOD2 specifically recognise PG-derived molecules of bacteria^39^. NOD1 recognises bacterial motifs containing the dipeptide or tripeptide-*meso*DAP or lanthionine^40^. NOD2 recognises the N-acetylmuramic acid (MurNAc) dipeptide present in almost all bacteria, as well as Mur-Tri-Lys present in some gram-positive bacteria^41^. Considering its key differences with bacterial PG, and notably the absence of MurNAc^42^, we investigated whether NOD receptors also recognise archaeal PG. However, none of the archaeal species were recognised by NOD receptors, regardless of whether they possess PG or not (Figure 3A and B).

**Figure 3.**
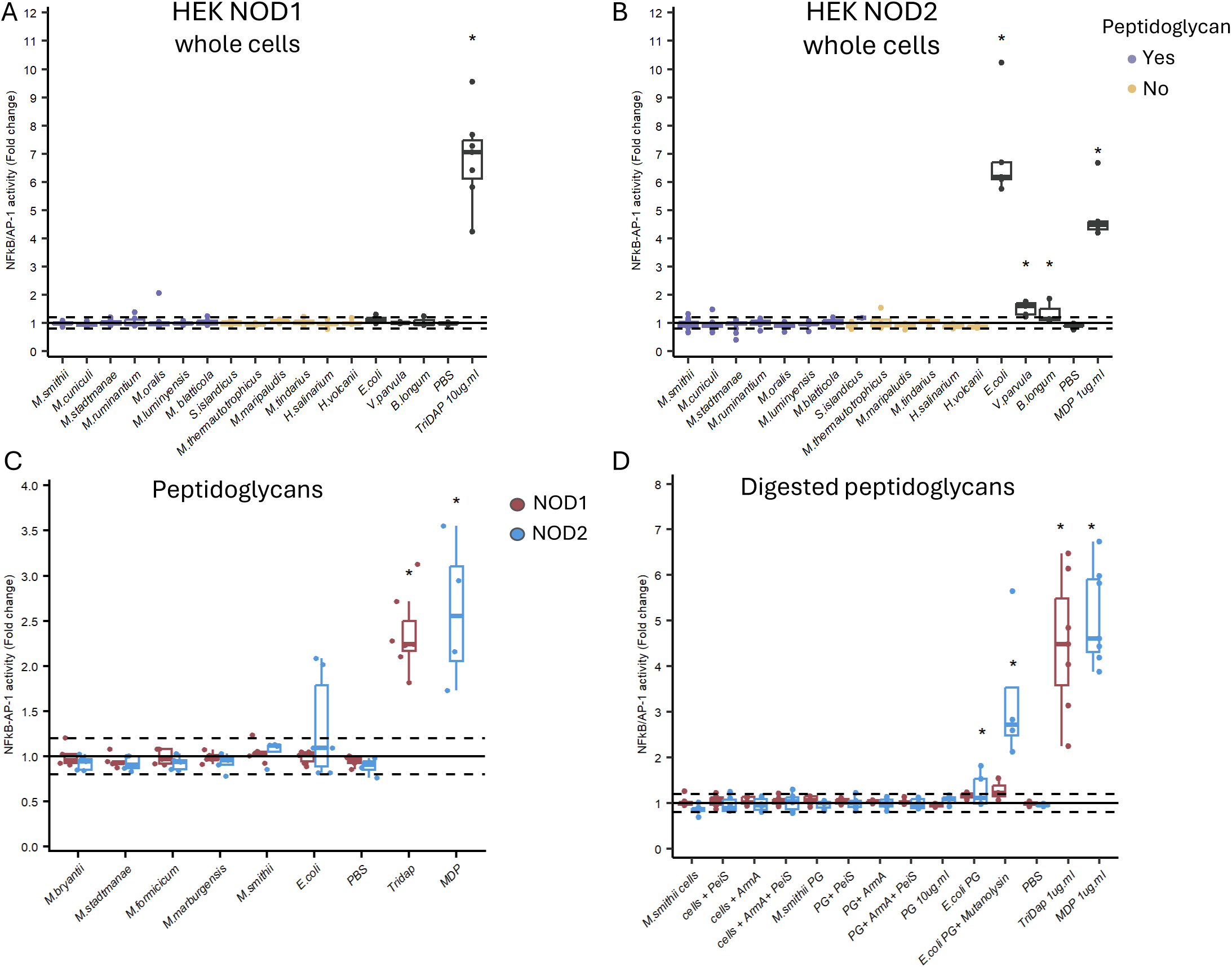
Archaeal PG is not a MAMP for NODs. NF-kB/AP1 activation upon exposure to archaeal cells on **A**) NOD1 and **B**) NOD2. **C**) NF-kB/AP1 activation upon exposure of cell lines to archaeal and bacteria purified PG. **D**) NF-kB/AP1 activation upon exposure of cell lines to archaeal and bacteria digested PG. Data are expressed as a fold change towards cells treated with PBS and represented as the median ± quartiles (N ≥ 3). *p value < 0.05 by Wilcoxon rank test.

Because in bacteria PG motifs are constantly released during cell division, we also tested the archaeal supernatants, but no activity was detected (Supplementary Figure 3A). The same lack of response was obtained using purified archaeal PG from five different species (Figure 3C). Undigested bacterial PG is a weak NOD inducer but its digestion by mutanolysin releases potent NOD activators. Thus, we digested both whole cells and purified PG from *M. smithii* with two hydrolytic enzymes having different cleaving site. However, neither digested *M. smithii* cells or purified PG resulted in NOD1 or NOD2 activation (Figure 3D).

### 4. Archaeal cells activate TLR2 by an unknown ligand

The bacterial ligands of TLR2 are mainly acylated lipoproteins and lipoteichoic acids^43^. Using whole archaeal cells, we show that 11 out of 15 species induced human TLR2, albeit with an overall lower activity compared to bacteria (Figure 4A). Halophiles had no activity on TLR2, which could be due to cell fixation. The variability of TLR2 activation may reflect biological variations, as also observed with bacteria. This activation was not restricted to the human receptor, as 6 archaeal species out of 10 showed activated murine TLR2 (Figure 4B). To test whether the innate immune activation observed in THP-1 cells (Figure 1C) was driven by TLR2, the receptor was blocked with a neutralising antibody prior to stimulation with archaeal cells. Compared to the control antibody, TLR2 blockage resulted in decreased NF-kB/AP-1 activation in THP-1 (Figure 4C).

**Figure 4.**
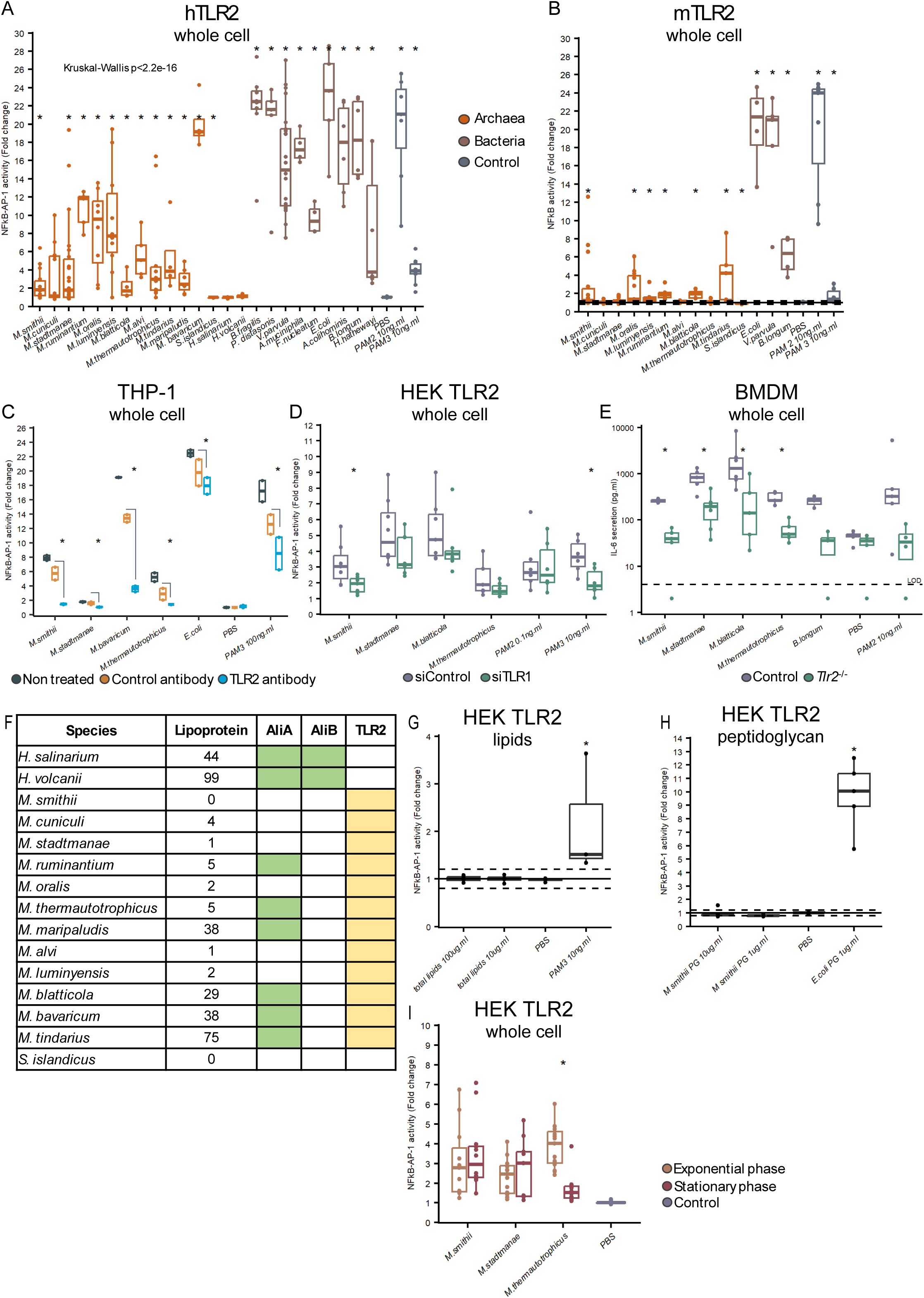
Archaea produce MAMPs recognised by TLR2. **A**) NF-kB/AP1 activation of hHEK TLR2 upon exposure to archaeal and bacterial cells. TLR2 activation difference between archaea and bacteria was assessed with a Kruskal-Wallis test. **B**) NF-kB activation of mHEK TLR2 cells upon exposure to archaeal and bacterial cells. **C**) NF-kB/AP1 activation upon exposure to archaeal cells of THP-1 cells pre-treated with TLR2 antibody, isotype antibody or non-treated. **D**) NF-kB/AP1 activation of HEK hTLR2 cells treated with siTLR1 and siControl. **E**) IL-c secretion of bone marrow-derived macrophages (BMDM) of Control and Tlr2^−/−^ mice treated with archaeal cells for 24h, limit of detection (LOD). **F**) Number of inferred lipoproteins in the genomes of the analysed archaeal species, presence of homologues of the lipidation genes AliA/B (green), and lack of correlation with TLR2 activity (yellow). **G-H**) NF-kB/AP1 activation of HEK hTLR2 cells after exposure to total lipids(G) or purified peptidoglycan (PG) (H) of M. smithii. **I**) NF-kB/AP1 activation of HEK hTLR2 cells after exposure to archaeal cells harvested during the exponential phase or the stationary phase. NF-KB activation results are expressed fold change compared to PBS treated cells, represented as the median ± quartiles (N ≥ 3). *p value < 0.05 by Wilcoxon rank test unless otherwise stated.

TLR2 can form homo– or heterodimers with TLR1 and TLR6, each having specific ligand affinity. Triacylated lipoproteins are the bacterial ligands of TLR2-TLR1, while diacylated lipoproteins are the ligands of TLR2-TLR6^44,45^. Our TLR2 cell line has a higher response to the synthetic ligands of TLR2-TLR6 (PAM2) than to the ligands of TLR2-TLR1 (PAM3), indicating that the TLR2-TLR6 dimer is more expressed than TLR2-TLR1 (Figure 4A). Therefore, the lower level of activation of archaea compared to the PAM2 control might indicate a higher affinity of the reporter for the TLR2-TLR1 heterodimer. To assess this hypothesis, we knocked down *TLR1* and *TLR6* in HEK TLR2 cells using siRNA. The results showed that TLR1 is involved in *M. smithii* activation, likely through the TLR1-TLR2 dimer, and a similar trend, although not statistically significant, was observed for the other archaea (Figure 4D, Supplementary Figure 4**A**). Interestingly, *M. stadtmanae* activity on TLR2 was decreased when TLR6 was knocked down (Supplementary Figure 4**B)**, indicating a potential role of the TLR6-TLR2 heterodimer in this species. These results suggest that archaea may produce different TLR2 ligands and involve different TLR2 dimers.

To assess our findings in a more complex and physiological model, bone marrow-derived macrophages (BMDM) of WT and *Tlr2^−/−^* mice were exposed to a selection of host-associated and environmental archaeal species and IL-6 secretion was quantified (Figure 4E). The four archaeal species induced IL-6 secretion, which was decreased in the absence of TLR2. This reduction was less pronounced in the case of *M. stadtmanae* and *M. blatticola*, suggesting the involvement of other PRRs. Our results are consistent with previous publications, and a recent pre-print showing that *M. stadtmanae* RNA is specifically recognised by TLR8 due to a specific enhanced endocytosis, which is not the case for *M. smithii*^46,47^.

Very little is known about lipoproteins in archaea^48^. Two archaeal enzymes (AliA and AliB) were recently proposed to be responsible for protein lipidation in *H. volcanii*^49^. These enzymes are analogues but not homologues of bacterial Lgt proteins. We analysed the presence of AliA and AliB homologues in our 15 archaea along with the number of putative lipoproteins as inferred from the presence of specific signal peptides (Figure 4F). As previously published, we only found AliB in halobacteria. In contrast, AliA homologues are present in 8 out of 15 species, but they exhibit patchy distribution, with no systematic presence within closely related taxonomic groups. Moreover, AliA presence does not correlate with TLR2 activity. We extracted *M. smithii* total lipids and tested if they activate TLR2 as well as TLR4, which also has lipid affinities, but no activity was observed on both receptors (Figure 4G and Supplementary Figure 4C). As lipoproteins can be associated to the PG in bacteria^50^, we tested the purified PG of *M. smithii* but again it did not activate TLR2, in contrast to the crude PG of *E. coli* (Figure 4H).

The fact that most archaea are recognised by TLR2 suggests either the existence of an alternative and yet unknown conserved pathway for lipoprotein synthesis, or that another type of triacylated compound acts as the archaeal TLR2 MAMP. To determine if the release of TLR2 activators depends on a specific growth phase, we compared the activity of cultures in exponential and stationary phases. No differences were observed between the two growth phases for both *M. smithii* and *M. stadtmanae*, but the activity was more pronounced in the exponential phase for *M. thermautotrophicus* (Figure 4I). In addition, multiple washes of the cells did not reduce overall TLR2 activity, as would be expected if the signal was due to molecules loosely attached to the cells (Supplementary Figure 4D).

## Discussion

Archaea are a stable component of multiple human microbiotas, from the skin to the oral and intestinal mucosa. However, our knowledge regarding their interaction with the host and role in human health remains very limited.

To our knowledge, this is the first study to examine MAMP-PRR interactions in a variety of archaea in terms of taxonomy, cell envelope structures and lifestyles. Our results show that archaea do not activate TLR4, TLR5, NOD1 and NOD2, the key receptors responsible for the sensing of bacterial cell envelope components. Specifically, we demonstrate a lack of activation by archaeal flagellin, PG, and lipids, whose bacterial counterparts are potent MAMPs. This result is intriguing and worth discussing. One possibility is that, during evolution, these receptors specialised on different components of the bacterial envelope (e.g., PG, lipoproteins, LPS, flagellin), and that the sequence or structure of their archaeal counterpart are too divergent to be recognised. The extent to which these receptors are flexible for different ligands is a matter of debate because of frequent contamination with LPS^51^. However, some TLRs, such as TLR2, are more promiscuous and have been suggested to participate in the signalling of additional types of MAMPs, such as the C-type lectins present in the envelope of fungi^48,49^.

We show indeed that archaea activate TLR2. These results contrast with previous work showing that *M. smithii* and *M. stadtmanae* cells activate monocyte-derived dendritic cells, but not TLRs or NODs, including TLR2 when expressed in HEK293 cells^24^. This discrepancy may be because the HEK293 cell lines used in that study do not overexpress NF-κB and are therefore less sensitive than the HEK blue recombinant cell lines that we used here and which overexpress NF-kB^52,53^. Moreover, we examined the activation of NF-kB whereas Bang et *al*. measured IL-8^24^. It is possible that the differences are also linked to unidentified different experimental conditions that could induce variation in the production and accumulation of TLR2 MAMPs in culture.

The archaeal ligand of TLR2 remains to be identified. Current knowledge of TLR2-TLR1/TLR6 indicates that these receptors can accommodate a ligand with or without a peptide part, although this is not common^54^. It may not be ruled out entirely that the observed TLR2 activity originates from exogenous ligands. However, we show that extensive washings of archaeal cells did not eliminate the activity. Furthermore, the different archaea grown in the same media do not signal in the same extent, which is difficult to reconcile with a common contaminant. Therefore, our result support the existence of archaeal TLR2 ligands which remain to be identified. Future efforts when genetic tools become available for a larger range of archaea, including host-associated members, will be key to identify the TLR2 agonist(s).

A recent *in vitro* study showed that the human and murine Mincle receptors (CLEC4E), belonging to the C-type lectin receptor (CLR) family, recognise archaeal synthetic glycolipids from *M. smithii* and *M. stadtmanae*^55^. Mincle has been shown to bind to fungal and human glycolipids, mycolic acids of Mycobacteria, the S-layer of Lactobacilli, and undetermined molecules of Leishmania ^56–59^. However, it remains to be determined whether archaeal cells expose glycolipids to Mincle and induce signalling. HEK cells and non-activated THP-1 cells do not express Mincle or its co-receptor, hampering the analysis of Mincle activation in a physiological context. Despite our attempts, we were unable to construct a fully functional reporter system of Mincle and Dectin-2 in HEK cells. The role of other lectin receptors has not yet been investigated, but preliminary tests indicate that archaea are not recognised by Dectin-1 (CLEC7A) (Supplementary Fig 5A). Interestingly, TLR2 have been shown to form heterodimers with CLRs, including Mincle^60^. TLR2 and Mincle have also been shown to act synergistically to detect Corynebacteria^61^. In the future, it will be interesting to investigate the role of CLRs and their potential synergy with TLR2 in the immune activity of archaea, notably in relation with the adjuvant properties of archaeal lipids^62,63^. Finally, if the cell envelope is largely not recognised by the innate immune system, other molecules may play an important role in the archaea-host interplay. A previous study and a recent pre-print study showed for instance that the RNA of *M. stadtmanae*, but not that of *M. smithii*, is specifically recognised by TLR8, in relation to the specific capacity of the cells of this species to be phagocytised by immune cells^46,47^.

Archaea have co-evolved with great apes and hominids as commensals for millions of years^64^. However, they do not meet the Koch postulate criteria for pathogenicity^11^. If, as largely assumed, the evolution of the immune system is driven by pathogenicity, one might therefore expect that archaea remain unrecognised. However, our and previous results show that archaeal MAMPs are recognised by at least three receptors (TLR2, TLR8, and Mincle). One hypothesis is that the presence of archaea at infection sites, where their molecules could encounter both innate and adaptive immune cells, may have been sufficient to select for PRR recognition, but not as strongly as bacterial and fungal pathogens.

Several gut-associated bacterial species show selection toward a reduction of their immunogenicity. For example, immunogenic flagellation is either counter-selected, resulting in loss or immunosilencing of flagellins with an antagonistic role on TLR5^38,65^. However, this scenario cannot explain our results since we do not detect a distinct immune recognition of archaea by TLR and NOD receptors depending on whether they are host-associated or not.

Another possibility is that commensal interactions between archaea and their host drove their detection by some PRRs. In fact, while the MAMP-PRR system has been discovered and investigated largely in the context of infections, an increasing number of studies suggests that these interactions play a role not only in immunity to pathogens but also in the regulation of host physiological processes by commensal microorganisms^66,67^. Under conditions of homeostasis, MAMPs cross in fact the mucosal barriers and interact with PRRs in distal organs, with deep impacts^66,68^. TLR2 is involved in various host homeostatic processes at the gut level^69^ as well as in the modulation of physiological functions in the brain, liver and angiogenesis^69–74^. This opens the question of whether PRR recognition is necessarily shaped by the history of infections, or if can also be driven by commensal interactions. If no pathogen exists among archaea, this makes them unique models to investigate the issue.

Finally, our results offer material to refine understanding of the evolutionary dynamics between PRR and MAMPs, as recently discussed^75^. A key question is how and why only specific molecules were selected for recognition as MAMPs by PRRs in mammals and other eukaryotes. Studies on the subject are however limited. Work on the immunogenicity of LPS in deep-sea bacteria has revealed that these bacteria exhibit much more variability in their lipid A, which in turn are less detected by TLR4, than the more conserved molecules in human-associated species ^76^. These results suggest that host pressure is a key element driving evolution of cell envelope MAMPs and the ability of PRRs to detect them. In contrast, our results show that this might not be the case of archaea, as we observe no difference in TLR2 recognition between host-associated and environmental species. This also further suggests that the MAMP recognized by TLR2 is a conserved and essential component of archaeal cell. Future development of key tools and murine models will open exciting avenues of research to understand the driving forces of PRR-MAMP selection and their impact on the interactions between humans and archaea.

## Material and Methods

### 1.1. Reagents

PAM2SK4, Pam3CSK4, MDP, TriDAP, Flagellin-ST ultra and PGN-EK were purchased from Invivogen. TNF-a (Peprotech, 300-01A). Ultrapure LPS *S. abortus* S-type was purchased from Enzo Life Science. Antibiotics used for the selection of the human and murine cells were purchased from Invivogen. Anti TLR2-IgA2 antibody (htlr2mab7-2) and Anti-β-Gal-hIgA2 (bgal-mab7) from Invivogen were resuspended in PBS.

### 1.2. Maintenance of cell lines

Raw 264.7 and THP-1 XBlue (Invivogen) cell lines were maintained in RPMI 1640 Glutamax (Gibco) supplemented with 10 % fetal bovine serum, 1 % non-essential amino acids (Gibco), 1 mM Sodium pyruvate (Gibco) and 10 mM HEPES (Gibco) at 37°C in a 5 % CO2 incubator. HEK-293 cells were cultured in DMEM Glutamax supplemented with 10 % fetal bovine serum (Gibco), 1 % MEM non-essential amino acids (Gibco) at 37 °C in a 5 % CO_2_ incubator. Serum was decomplemented by heat inactivation at 56 °C for 30 min. Mycoplasma and bacterial contaminations were tested regularly by PCR as previously described^77^ and using HEK TLR2 cells. The cell viability was monitored by MTS measurement using the CellTiter 96 Aqueous One solution (Promega) according to the manufacturer’s recommendations.

### 1.3. Reporter system cells

Commercially available NF-kB-AP1 reporter from Invivogen were used: HEK-293 Null1, TLR2, TLR4, NOD1 and NOD2, THP-1 XBlue. HEK TLR5 were construct as stable cell lines, overexpressing human TLR5 and murine TLR5 were generated by transfecting 10 µg of the pUNO1-hTLR5 or mTLR5 (Invivogen) in HEK Null KD TLR5 background with 15 µl of Lipofectamine 3000 and 20 µl of Reagent. Stable colonies were selected and cultured in the presence of blasticidin (Invivogen) for at least five weeks. The colonies were then tested for activation in response to flagellin.

Raw 264.7 NF-kB were constructed as stable cell line by plating cells at 0.25 x 10^+6^ cells in 6 well plates. The next day cells transfected with 1 µg of pNIFTY2 plasmid mixed with 5 µl of Reagent in 200 µl of Buffer using the Jetprime transfection (PolyPlus). After 24 h, fresh RPMI was added with Zeocin, and a pool of stably transfected cells were selected from 4 weeks. The NF-kB reporter system was validated by testing the cell response to LPS and PAM2 and PAM3.

For each experiment, HEK reporter cells were seeded at 30 000 cells per well, Raw were seeded at 20 000 cells/well and THP-1 XBlue at 80,000 cells/well in a 96 well plate 24 h prior stimulation in technical duplicate. After 24 h of incubation, 20 μl of supernatant was added to 180 μl of Quanti-Blue substrate (Invivogen) and read at 655 nM using a plate reader. Each sample was assessed in duplicate, and all experiments were tested in at least three biological replicates.

### 1.4. siRNA Knockdown

HEK cells were seeded at 10,000 cells per well in a 96 wells plate in 80µl per well and at 0.5 x 10^6^ cells in 6 wells plate for qPCR. The next day siRNA transfection was prepared by mixing 5 μM of siRNA control (ON-TARGETplus Nontargeting Control Pool D-001810-10-05), siRNA TLR1 (7096 ON-TARGETplus Human – SMARTpool L-008086-00-0005), siRNA TLR6 (10333 ON-TARGETplus Human, SMARTpool L-005156-00-0005) and 0,2 μl well DharmaFECT 1 in Optimem for a total volume of 20 µl per well, using DMEM without FBS. For the 6 wells plate, 5µM of siRNA were mixed with 10 μl of DharmaFECT per well in a total volume of 400µl. siRNA and transfection reagent diluted separately for 5 min in DMEM and then mixed and left for 20 min incubation at room temperature before addition the plate. The same procedure was repeated the day after for a second round of transfection. Before adding the new transfection volume, 20 µl were removed from each well. On the fourth day, cells were stimulated in duplicate for 24 h before revelation of the NF-kB activation with Quanti-Blue as described in section 1.3. Cells were recovered from the 6 well plates in 350 µl of RLT buffer with b-mercaptoethanol and 350% of 70% ethanol before freezing at –80°C until extraction.

### 1.5. RT-qPCR

Frozen cells were extracted using the RNeasy Mini kit (Qiagen) according to the supplier’s instructions. RNAs were eluted in 42µl and 2µg were used to perform reverse transcription using the using the High-Capacity cDNA Reverse Transcription Kit (Applied Biosystems). 10ng of cDNA were used per reaction using the Taqman Universal Master Mix II (Applied Biosystems) and 0.5 µl of probe in a total volume of 10µl on a CFX BioRad System. The following Taqman Gene expression assay probes were used: *TLR1* (Hs00413978_m), *TLR6* (Hs04975840_m), *TLR2* (Hs00610101_m) and *B2M* (Hs00187842_m). *B2M* was used for normalisation, and the fold change was calculated with the 2-ΔΔCτ method toward cells treated with the siControl. Samples were tested in experimental duplicates and at least in biological triplicates.

### 1.6. Neutralising antibody

For the neutralisation assay, THP-1 XBlue cells were seeded at 80,000 cells/well in 90µl. The next day, 10 µg.ml TLR2 antibody or Control antibody were pre-incubated for 1h at 37°C before stimulation with archaeal cells or controls for a total of 24h. NF-kB activation was assessed by Quanti-Blue as described above.

### 1.7. Bone marrow-derived macrophages

All animal experiments were approved by the committee on animal experimentation of the Institut Pasteur and by the French Ministry of Research (APAFIS #47193-2024020111556371 v2). Bone marrow-derived macrophages (BMDMs) were obtained after cervical dislocation of C57BL/6 mice of 5 weeks minimum, females and males. All the mice were bred by the Institut Pasteur. The control mice were Lgr5-GFP. After isolation of the femurs and tibias, the head of the bones were cut, and the bone marrow was flushed out in 150µl of PBS by a short centrifugation at 12 000 g. The cells were centrifuged at 4 °C and erythrocytes were lysed at room temperature (AKC buffer, Gibco A10492-01). Cells were then washed and counted before freezing in RPMI complete medium with the addition of 5% DMSO.

Cells were thaw and plated in 10cm petri dishes in 10 mL of differentiation medium: RPMI 1694 with 10% V/V heat-inactivated FBS (Gibco), 1mM sodium pyruvate (Gibco) and 1 X non-essential amino acids (Gibco), 1mM HEPES, 1 X antibiotic-antimycotic (Gibco) and 10 % V/V L929 cells supernatant to differentiate them to macrophages. L929 was obtained from confluent cultures in DMEM and filtered through 0,22 μm before freezing. Cells were incubated at 37°C, 5% CO2 and after 3 days, 3ml of differentiation medium was added. At day 6 days BMDMs were recovered by scraping in Cell Dissociation Buffer (Gibco). Cells were washed and counted to plated 0.1 x 10^6^ cells per well in 100µl in a 96 wells plate. At day 7 cells were stimulated for 24h with archaeal cells and controls. The supernatants were recovered the next day and frozen at –20°C until analysis.

### 1.8. ELISA

THP-1 XBlue cells were seeded at 1 x 10^6^ cells per well in 24 wells plate and stimulated the next day in a total volume of 400 µl. Archaeal cells at OD1 were added at 10 % v/v and incubated for 24h. The next day supernatants were harvest by centrifugation at 400 g for 10 min at 4°C. and transferred to clean tubes and frozen until use at –20°C. IL-6 and TNF were assessed using the ELISA Max Deluxe set kits (Biolegend) using half-area plates (3695 Costar) in a total volume of 50 µl. Absorbance at 450 nm was subtracted with the absorbance at 570 nm (Cytation). When IL-6 was measured below the limit of detection the minimum value (4 pg.ml for hTLR6 and 2pg.ml for mIL6) was set to perform statistical comparison. IL-6 quantification in the supernatant of BMDM cells was performed similarly.

### 1.9. Culture of archaea and bacteria

*Methanococcus maripaludis* S2 strain was obtained from Dr de Dios Mateos (Aalto University, Finland), *Haloferax* volcani H26 strain was obtained from Dr Lestini (Polytechnique, France) and *Halobacterium salinarium* FLX37 strain was obtained from Dr Marwan (University of Magdeburg, Germany). *Sulfolobus islandicus* REY15A cultures were obtained from are a kind gift of Diana Baquero Uriza (Institut Pasteur). Species information and culture media are listed in Supplementary Table 1 and Table 2. Cultures were harvested at late stationary phase to achieve the maximum biomass and centrifuged for 20 min at 3200*g* at room temperature. Non inoculated culture media and supernatants were filtered on 0.22 µm PES Millex filers (Millipore). Cells were washed with 10 ml PBS, centrifuge 20 min at 3000 *g* and resuspended in PBS at OD=1 and kept at –20°C until test.

Absolute cell number of *M. smithii* and *M. stadtamane* cells was determined from cell washed with PBS and stained with in Truecount tubes containing beads (BD 663028) and counted by flowcytometry with an Astrios cell sorter (Beckman Coulter). OD=1 correspond to ≈ 9 x 10^+8^ *M. smithii,* 4 x 10^+8^ *M. stadtamane,* 1.1 x 10^+9^ *M. luminyensis, 1.2 x 10^+S^ Haloferax volcanii*^78^, 5 x 10*^+8^ M. thermautotrophicus* cells/ml and to 1.5 x 10^+9^ CF for *H. salinarium*^79^. In HEK 96 wells assay 10µl of archaeal cells at OD1 were applied per well, corresponding to ≈0.4-1 x 10^+7^ archaeal cells for 10^+5^ human cells (MOI 40-100).

Cultures of halophilic archaea were harvested in stationary phase, and the supernatant was filtered at 0.22 µm after 20 min centrifugation at 3200 *g*. Cells were resuspended in 300 µl medium and 100 µl of 37 % PFA was added for fixation, 45 min at 37°C. Cells were then washed twice with 1 ml PBS and 10 min centrifugation at 5000 *g* and kept at –20°C until test.

*E. coli* Nissle 1917 was grown in LB, *V. parvula* SKV38 in SK lactate medium and *B. longum* (CIP 6462/DSM 20219) in MRS for 24 h. Cultures were harvested after 24 h and centrifuged for 20 min at 3200 *g* at room temperature. Non inoculated culture media and supernatants were filtered on 0.22 µm PES Millex filers (Millipore). Cells were washed with 10ml PBS, centrifuge 20 min at 3000 *g* and resuspended in PBS at OD=0.5 and kept at –20°C until test. To prevent *E. coli* growth during tests, cells were killed by heating at 99°C for 20 min before freezing for further use.

### 1.10. Electron microscopy

Fresh cultures of were blotted in carbon-coated (200 mesh) glow discharged copper grids (2 mA and 1.5–1.8 × 10^−1^ mbar for 1 min in a glow discharge system (ELMO, Corduan) and negatively stained using 2 % uranyl acetate (wt/vol). Images were acquired using a Tecnai 20 equipped with a field emission gun and operated at 200 kV (Thermo Fisher Scientific).

### 1.11. Flagella extraction and deglycosylation

Flagella of halobacteria were extracted from 200 – 250 ml of stationary cultures. After centrifugation at 3000*g* for 20 min, the supernatant fractions were concentrated by centrifugation at 16 000*g* at 4°C on 3 kDa Vivaspin columns (Satorius).

Before deglycosylation, the concentrated material was lyophilised resuspended in 300 µl of Trifluoromethanesulfonic acid (TFOH, Sigma) in a dry-ice cold bath and incubated for 3 h with gentle stirring. The reaction was neutralised with 60 % pyridine and using paper pH tests. The deglycosylated flagella were subsequently washed 3 times with 500µl PBS using 3 kDa Amicon Ultra columns and resuspended in the original volume. Each sample was divided in two, with half been sonicated 2 x 45 s at 30 % amplitude on ice. Samples were kept at –20°c until use. Samples were diluted 1/10 before test to avoid the death of HEK cells.

### 1.12. Archaeal peptidoglycan extraction

Archaeal cells were pelleted by centrifugation at 4000 *g* for 20 min. Cell pellets were resuspended in 10 ml of PBS and added dropwise to 10 ml of 8 % w/v SDS preheated in a boiling water bath and incubated at 100° C for 1 h. The cell lysate was cooled to room temperature and sacculi were pelleted by ultracentrifugation at ≈150 000 *g* for 1 h. The sacculi were resuspended in 8 ml of ddH_2_O and washed a minimum of 10 times by centrifugation at 17,000 *g* to remove residual SDS. The pellets were then incubated at 37°C in 8 ml 50 mM Tris-HCl pH 7.0 with 100 μg.ml alpha-amylase for 2 h with constant agitation. Then 10µl.ml of benzonase 25 U.µl (Novagen) was added, and the digestion was incubated for another 2 h at 37°C. Finally, 100 µg.ml of trypsin was added, and the digestion was incubated overnight at 37°C with constant agitation. SDS was added at a final concentration of 1% to inactivate the trypsin at 95°C for 15 min. Sacculi were washed in ddH_2_O 4 times to remove the SDS and resuspended in 8 ml 8 M LiCl and incubated at 37°C with constant agitation for 30 min. Pellet was pelleted by centrifugation at 17,000 *g* for 10 min and resuspended in 100 mM EDTA to chelate and remove remaining LiCl along with other ions and incubated at 37°C with constant agitation for 15 min. Sacculi were pelleted by centrifugation at 17,000 *g* for 10 min and washed twice in 8 ml ddH_2_O. Finally, the pellet was washed with 1 ml of 100 % acetone and the major aggregates removed by short sonications. The pellet was washed twice with ddH_2_O and lyophilized by freeze drying.

### 1.13. Purification of archaeal peptidoglycan hydrolytic enzymes

*E. coli* BL21 cells containing plasmids expressing C-terminally 6 x His tagged recombinant ArmA were cultured overnight at 37°C in LB supplemented with 100 μg.ml ampicillin. Overnight culture was diluted 1/1000 in autoinduction medium as described previously (in Smith et al. in revision) and cultured at 37°C, shaking at 180 rpm until OD_600_ 0.7. Cultures were transferred to 30°C and grown overnight. Cells were centrifuged at 6000 g for 20 min, resuspended in buffer A (50 mM Tris-HCl pH 7.5, 300 mM NaCl, 15 mM imidazole, 5 % v/v Glycerol) supplemented with protease inhibitors (Roche cOmplete EDTA free) and benzonase (Novagen). Cell lysis was achieved using a cell disruptor (Constant Systems) at 1.5 kbar pressure. Cell debris was removed by high-speed centrifugation at 50 000x *g* for 1 h at 4°C. The cell lysate was loaded onto a Co-NTA affinity column (HiTrap talon 5 ml GE healthcare) at 10°C. The protein was then eluted using a linear gradient of buffer B (50 mM Tris-HCl pH 7.5, 300 mM NaCl, 500 mM imidazole, 5 % glycerol). Proteins were further purified by size exclusion chromatography (SEC) using a Superdex 200 16/60 column and SEC buffer (20 mM Tris-HCl pH 7.5, 300 mM NaCl). Proteins were stored at –20°C.

### 1.14. Digestion of archaeal peptidoglycan

Either 10 µg of archaeal *peptidoglycan* or 1.25 mg of *M. smithii* cells were incubated with of ≈13 µg PeiS (2.63 mg.ml), 8 µg of ArmA_cter_ or control buffer (pH 7.5, 50 mM HEPES, 160 mM NaCl, 2 mM CaCl2, 2 mM MgCl_2_ with 1 mM DTT freshly added). The reaction was incubated at 37°C in anaerobic conditions for 24h. At the end of the reaction, 50µl of PBS was added and cleavage products were recovered by filtration on Amicon Ultra-0.5 3 kDa filter with 2 centrifugations of 30 min at 14 000x *g* and 4°C. As control, 0.5 mg.ml of *E. coli* peptidoglycan (Invivogen PGN-EK) was digested in a 20 mM Tris 150 mM NaCl pH 7.5 Buffer, with 40 U of Mutanolysin (Sigma M9901) per 100 µg of peptidoglycan, overnight at 37°c with agitation. The bottom fraction <3kDa were frozen and the activity assessed on human cells.

### 1.15. Total lipid extraction

*M. smithii* was grown in the same medium as described in SUP Table 1 but a fermenter was used to obtain a larger biomass. The extraction process began with 1 g of cells, which were washed twice with ddH_2_O at 400 *g* for 5 min to remove any culture debris. The cells were spun again at 3200 *g* for 20 min, washed with 50 ml of 0.9 % NaCl 0.02 % Triton X-100, and rinsed twice with 50 ml of 0.9 % NaCl before lyophilisation using a freeze dryer. The cells were transferred to a 25 ml Corex glass tube and layered with 2.5 ml of CHCl₃, followed by 5 ml of MeOH and 2 ml of 0.3 % NaCl. The tubes were mixed and incubated at 80 °C for 15 min, followed by an hour-long incubation at room temperature on a vortex. The tubes were spun at 4000 rpm for 15 min at room temperature, resulting in a white pellet and clear supernatant. The resulting supernatant was transferred to a clean Corex tube, resuspended in 2.5 ml of CHCl₃, 5 ml of methanol and 2 ml of 0.3 % NaCl, and vortexed for 1 h at room temperature. After centrifugation at 4000 rpm for 15 min, the supernatant was transferred to a new Corex tube, to which 2.5 ml of CHCl₃ and 2.5 ml of NaCl were added and mixed. After 15 min of centrifugation at 4000 rpm, the upper fraction and the white interphase were removed, and the bottom organic phase retained. The organic phase was purified again by adding 3 ml of 0.3 % NaCl and spinning for a further 7 min at 4000 rpm. The bottom organic phase, which contained the total lipids, was recovered and placed in pre-weighted vials to evaporate the CHCl₃ under a N₂ stream. The lipids were kept lyophilised at –20 °C until use. To test the lipids on HEK cells, 500 µg of lipids were resuspended in 500 µl of PBS, vortexed for 2 x 20 s to form liposomes, and kept at 4 °C until use.

### 1.16. Bioinformatic analyses

All proteins encoded in the 15 archaeal genomes (30,546 proteins), listed in Supplementary Table 3, were scanned to identify secretion signals using SignalP 6.0 ^80^. SignalP 6.0 identifies five types of signal peptides, including distinct predictions for lipoproteins secreted via Tat (TATLIPO) and Sec (LIPO) pathways. We considered only positive predictions with a probability ≥80% for all types of signal peptides.

To identify potential homologs of AliA (arCOG02177) and AliB (arCOG02178), we conducted HMM-based homology searches using profiles obtained from the online arCOG v14 database (https://ftp.ncbi.nlm.nih.gov/pub/wolf/COGs/arCOG/) using HMMSEARCH ^81^ with an e-value threshold of 1×10^−3^. The resulting homologues were aligned using MAFFT v7.407^82^ with the L-INS-I option. Alignments were manually curated to remove non-homologous sequences and used to build more specific HMM profiles to carry out further iterative searches until no additional homologue was found.

To search the D0-D1 flagellin motifs, the proteomes of the 15 archaeal species were screened for the presence of the following motifs from different bacteria:

LALLKAN, FMPLED, KSSRFSKDRNLYVND, KDLPKDLQKKVLAKESVRVYL, LQRVR*L*VQ,LQRIRELA VQA, LDTVKVKATQ

### 1.17. Statistical Analysis and Graphics

Statistical analysis was done using non-parametric Wilcoxon rank test, otherwise the specific test is mentioned. Graphical representations show single data points using ggplot boxplot, resenting the median, the first and third quartiles. Data come from at least three independent biological experimentations. Data were analysed using R 4.5.2 and RStudio 2023.06.0 B 421 software. Graphics and statistical analysis were produced with ggplot2, ggsci, gridExtra, ggsignif, and ggpubr packages. Adobe Illustrator 29.8.4 was used to generate figures. Data are expressed as fold change toward cells treated with PBS or the corresponding control media otherwise stated. The absence of symbol indicates a non-significant p-value.

## Author contributions

CMG conceived the study, performed the experiments and analysed the data. CG helped with the BMDM experiments. BB performed the bioinformatics analyses. CW advised and provided material for some experiments. SG provided infrastructure and funding. CMG and SG wrote the paper, with input from GB and CW.

## Acknowledgments

We thank Robert Smith for help with archaeal PG extraction and Olivier Danot for help with purification of archaeal PG hydrolytic enzymes. We are grateful to Diana Baquero Uriza for providing the *S. islandicus* cultures and help on other archaeal cultures. We thank Leticia Beltran for transmission electronic microscopy of archaeal flagella. We are grateful to Arnaud Chastanet for help with *M. smithii* lipid extraction and to Johan Cornil for help in the deglycosylation protocol of archaeal flagella.

## Data availability statement

All data are contained in the article and the supporting information.

## Open Access

This article is licensed under a Creative Commons Attribution 4.0 International License CC BY-NC-ND 4.0, which permits use, sharing and reproduction in any medium or format, as long as you give appropriate credit to the original author(s) and the source, provide a link to the Creative Commons licence, and indicate if changes were made. The images or other third-party material in this article are included in the article’s Creative Commons licence, unless indicated otherwise in a credit line to the material. If material is not included in the article’s Creative Commons licence and your intended use is not permitted by statutory regulation or exceeds the permitted use, you will need to obtain permission directly from the copyright holder. To view a copy of this licence, visit https://creativecommons.org/licenses/by-nc/4.0/legalcode Opens in new window.

## Supplementary Figures

**Supplementary Figure 1**. **A)** NF-kB/AP1 activation level after 24h exposure to archaeal cells on HEK Null cells. **B)** NF-kB/AP1 activation level after 24h exposure to archaeal cells on TLR4 HEK cells. Results are expressed fold change compared to PBS treated cells, represented as the median ± quartiles of fold change toward cells treated with PBS (N *≥ 3). *p value < 0.05 by Wilcoxon rank test. Blue: Host-associated; Orange: Environmental species*.

**Supplementary Figure 2**. Purified flagella obtained from H. salinarium were negatively stained and imaged by transmission electron microscopy.

**Supplementary Figure 3**. Archaeal supernatants do not trigger NODs. NF-kB/AP1 activation exposure of NOD1 and NOD2 cell lines to archaeal supernatants. Results are expressed as fold change towards cells treated with control media and represented as the median ± quartiles (N ≥ 3). *p value < 0.05 by Wilcoxon rank test.

**Supplementary Figure 4**. **A**) mRNA levels of TLR1 and TLR6 upon treatment with the siRNA targeting siTLR1, siTLRc or siControl (ΔΔCt toward siControl. **B**) NF-kB/AP1 activation of HEK hTLR2 cells treated with siTLR6 and siControl **C**) M. smithii lipids do not trigger TLR4. NF-kB/AP1 activation following exposition of HEK TLR4 to different concentration of M. smithii total lipids. Results are expressed as fold change towards cells treated with PBS and represented as the median ± quartiles (N =4, except for the LPS condition N=2). **D**) NF-kB/AP1 activation following exposition of HEK TLR2 to cell pellet washed 1, 2 or 3 times or control media with PBS. *p value < 0.05 using Wilcoxon rank test.

**Supplementary Figure 5**. NF-kB activation following exposition of HEK Dectin-1 archaeal cells. Results are expressed as fold change towards cells treated with PBS and represented as the median ± quartiles. *p value < 0.05 by Wilcoxon rank test.

