## Supplementary figures and images for "Archaeal cell envelopes selectively activate innate immunity through TLR2 in contrasts with multi-receptor bacterial sensing"

### Supplementary figures

**Supplementary Figure 1**

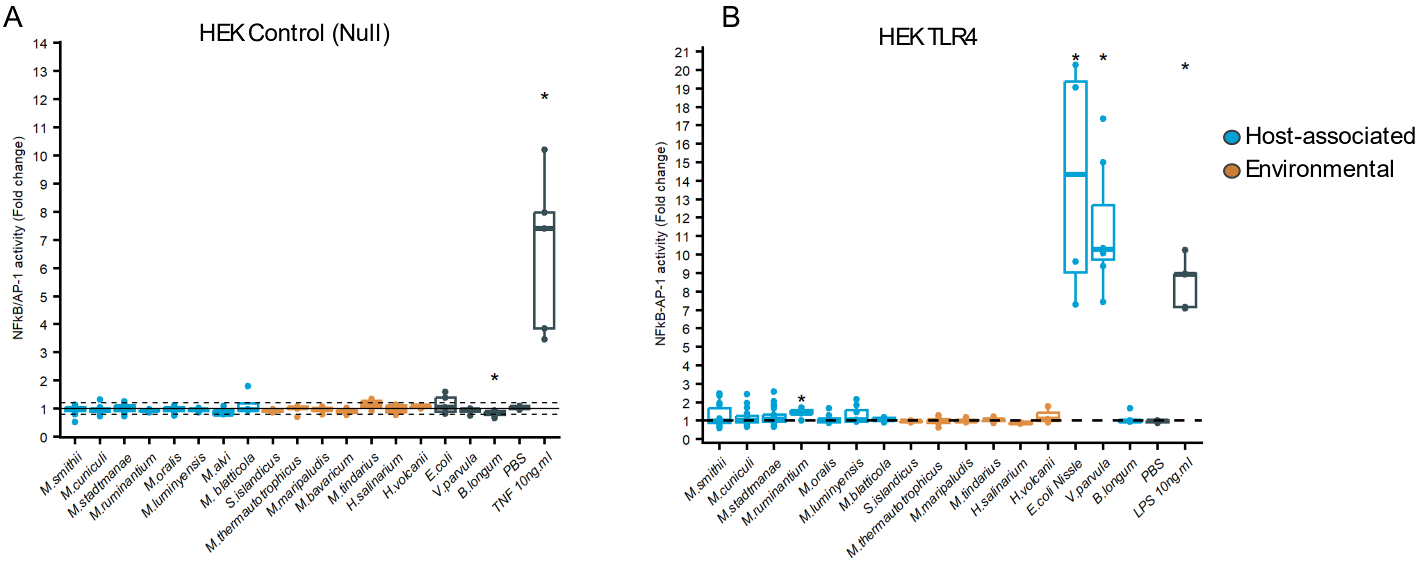

**Supplementary Figure 2**

A

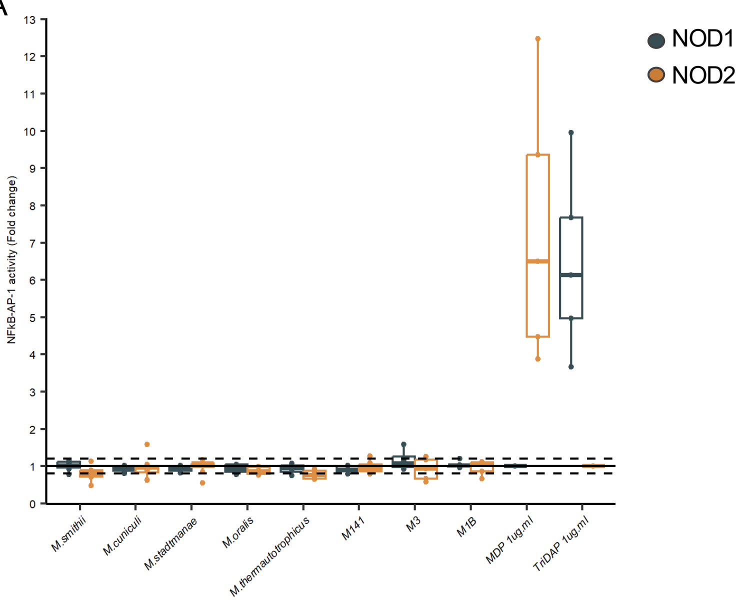

**Supplementary Figure 4**

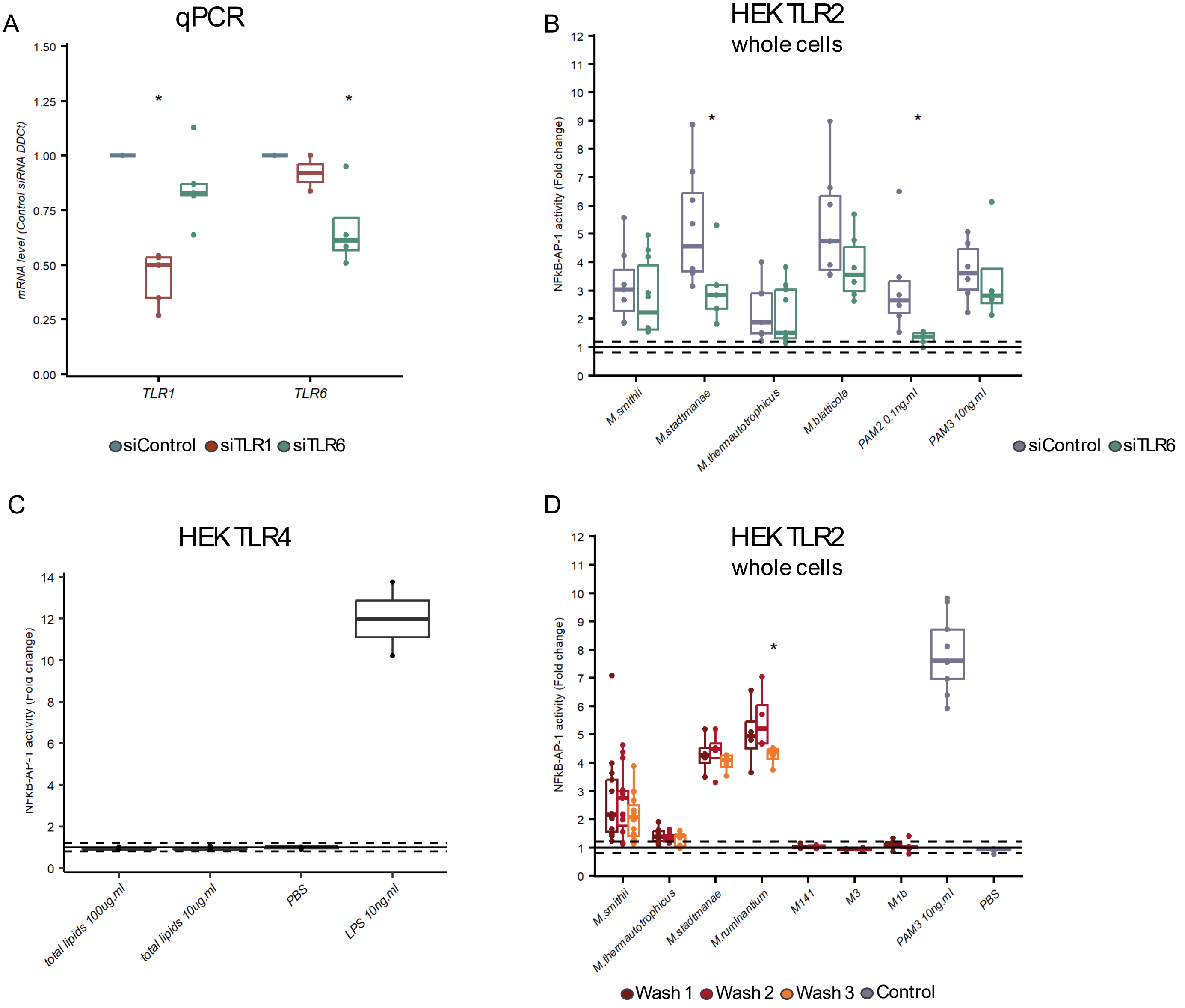

**Supplementary Figure 5**

A

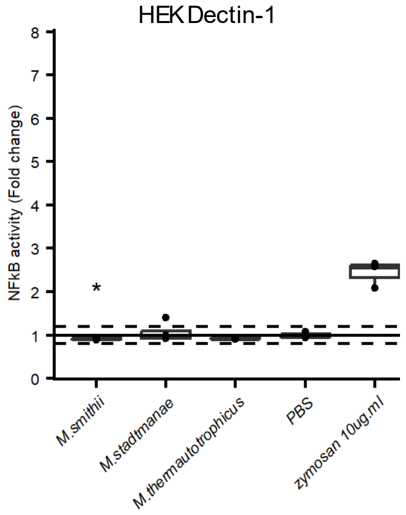
